# A target-free approach to estimate thermal camera pose in LiDAR scenes of feature-deficient environments

**DOI:** 10.1101/2023.10.09.561532

**Authors:** Julian Jandeleit, Thejasvi Beleyur, Bastian Goldlüecke

## Abstract

Modern animal behaviourists are able to collect vast amounts of data from multiple sensors. Sensor fusion however still remains a challenge given the unique sensor types biologists use. Here we study thermal camera-LiDAR alignment. While LiDAR-RGB scene alignment is well-established in feature-rich scenes with reliable inter-camera correspondences, thermal scenes are often captured with low-resolution devices and suffer from halo and history effects, and thus require sensor-specific algorithms. To deal with this, we present the depth-map correspondence (DMCP) algorithm. DMCP is a semi-automatic algorithm to fuse LiDAR with feature-deficient thermal scenes without explicit calibration objects. The user annotates at least 4 corresponding points on the thermal image and a depth map of the LiDAR scene, which allows us to compute the position and orientation of the thermal camera within the scene. We quantify the accuracy of alignment using 2D reprojection error and 3D nearest-neighbour distances of known world points across experimental trials. A median reprojection error of 7.7 (95%ile: 3.7-23.5) pixels was achieved. Known objects lying on the LiDAR mesh showed a median nearest-neighbour mesh distance of 0.12 m (95%ile: 0.007-5.61 m). We present the first such alignment of feature-deficient thermal scenes and LiDAR data. DMCP has two advantages 1) it is computationally straightforward to implement and 2) requires minimal user annotation to work. DMCP could be broadly applied to any type of mesh-image alignment problem. Finally we provide DMCP and the Ushichka dataset as a baseline for further research on the challenging problem of subterranean thermal-LiDAR alignment.

## 1 Introduction

Animal behaviorists today are able to collect types of data from multiple sensors such as thermal video and even LiDAR scans of the natural environment. Contextualising the animal’s behaviour however requires bringing the multi-sensor data to a common coordinate-system. Animals occupy a wide set of physical environments (aquatic, terrestrial, subterranean), each of which constrain sensor performance (e.g. spectral content and contrast). However, state-of-the art methods to handle and align sensor data are typically centred around human and industry environments (e.g. artificial and urban settings) with their own unique features. State-of-the-art methods may thus fail when used with unconventional data from natural environments, necessitating the development of new algorithms.

In particular, the *Ushichka* dataset [2] which we are interested in is a multisensor dataset with multi-channel audio, multi-camera thermal video and LiDAR scans of the Orlova Chuka cave system in Bulgaria. Echolocating bats live in the cave, and fly around the recording volume across the night. Their flight behaviour and echolocation in groups is of particular interest to sensory biologists and collective behaviorists. Each of the sensors in *Ushichka* captures an important aspect of the animals’ behaviour, but a contextual understanding is achieved only by aligning the three sensors into a common coordinate system. To understand a bat’s flight decisions we need to know its flight path with respect to the cave’s walls. Aligning the LiDAR and the camera coordinate systems allows this contextual understanding. Aligning 3D meshes and thermal cameras is typically done with calibration objects in the scene [18,11]. The *Ushichka* scene however does not contain a calibration object visible to both LiDAR and thermal cameras. We thus need to make use of the naturally available features in the data.

While methods using environmental features developed for visible light camera and LiDAR alignment without calibration objects exist [14,17], all of our alignment attempts based on feature and photo-consistency based methods from previous works such as SIFT, LGHD and space-carving [15,1,13] approaches failed to provide enough scene points necessary for alignment [8]. A main reason for the failure of these methods on the *Ushichka* dataset is likely the unique nature of the subterranean thermal scenes, which exhibit extremely low contrast across the scene. The cave system holds a fairly stable temperature of around 10^*°*^C all year round, and shows very little variation in temperature from one part to another (*pers. obs*.). The thermal scene is also very self-similar. The rocks in the scene are difficult to uniquely identify, the smooth walls dominate portions of the camera view - resulting in features with low descriptive power.

We present the semi-automated Depth-Map Correspondence (*DMCP*) algorithm developed specifically to handle the image-mesh alignment of feature deficient scenes such as those in *Ushichka*’s thermal data (See Fig. 1). *DMCP* consists of the following steps:

**Fig. 1:**
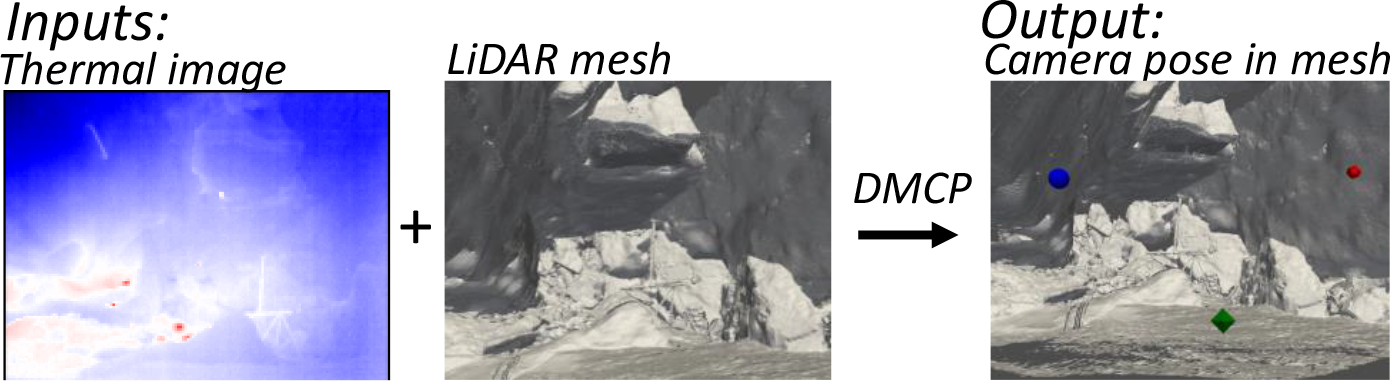
Schematic showing the required raw data inputs and resulting output from the depth map correspondence (DMCP) algorithm. While designed for thermal camera pose estimation, DMCP can work for any type of image-pointcloud or image-mesh sensor data.

1. The user chooses a view of the mesh interactively to approximately match the thermal image. A depth map of the mesh is then generated from the viewpoint. A depth map is image-like and thus eases comparison with the thermal image.
2. The user annotates a minimum of 4 point-to-point correspondences between the thermal image and the depth map (Fig. 2). Each 2D point in the depth map corresponds to a 3D point in the LiDAR scene.

**Fig. 2:**
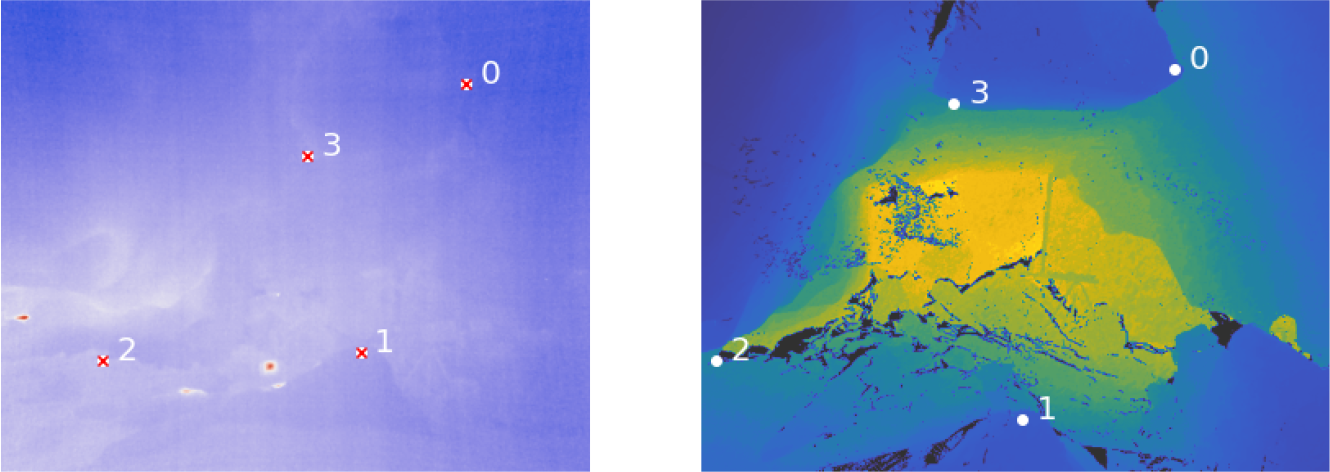
Example image and depth map from *Ushichka* experiment. White points indicate annotated pixels. A red *x* indicates the reprojected depth map correspondence.
3. The extrinsic pose of the camera in the LiDAR scene is estimated from the user-annotated correspondences.
4. The transformation matrix that converts 3D points from camera world space to LiDAR space is calculated. This is done by passing points through camera space and solving the absolute orientation problem.

## 2 Methodology

### 2.1 The *DMCP* algorithm

Most of the terminology and mathematical background is used as defined in [6]. The extrinsic matrix *E* is the transformation that brings points in world space to camera space. The camera intrinsic matrix *K* projects points from camera space into pixel space. Pixel space describes the coordinates of points that lie on the image itself. The pinhole projection matrix is denoted as *P* = *KE*. It projects points from world space to pixel space. Each extrinsic matrix *E* has a respective pose matrix *C*, which is the inverse of *E* when expressed in homogeneous coordinates. Objects in homogenous coordinates are denoted with a hat (e.g *Ê*).

We call the coordinates the LiDAR mesh is expressed in the world space. The camera to be aligned is given in its own associated space defined by its extrinsic matrix. To not confuse it with the world space given by the mesh, this space is called the camera’s native space. In the case of the *Ushichka* dataset, the extrinsic matrix can not be normalized to the identity as there are three thermal cameras calibrated relative to each other extrinsically so that they share their native space. To distinguish coordinate systems, the system referred to is denoted in the subscript of objects. For example *P*_*native*_ denotes the pinhole projection matrix that projects points from native space to pixel space.

Given a thermal image *I*_*native*_ captured by a camera *P*_*native*_ = *K*_*native*_*E*_*native*_ in native space, the task is now to find a projective transformation *M* from native to world space such that the coordinates of a 3D point *A* in the two systems are related by the affine transformation *Â*_*world*_ = *M Â*_*native*_.

A concise overview of the *DMCP* Algorithm is presented in pseudocode here. Algorithm 1 shows the complete workflow used to register native and world space. It makes use of the Sparse Camera Algorithm (SCA, Algorithm 2), which includes the main operations necessary to find the transformation. The mathematical details are laid out in the paragraphs below.

### Point Annotation

The user first chooses a view point to approximately match the captured thermal image *I*_*native*_. Intrinsics are typically chosen to be the same as for the thermal image, while the pose is selected by moving through an interactive rendering of the scene. From the selected view point, we render a depth map *I*_*dm*_ of the mesh with [16] according to the chosen projection matrix *P*_*dm*_. The pose *C*_*dm*_ = *R*_*dm*_ *T*_*dm*_ of the virtual camera that captured the depth map is saved. The user now annotates at least 4 corresponding points on *I*_*dm*_ and *I*_*native*_. We denote the two corresponding sets of points by *cps*_*dm*_ and *cps*_*image*_, respectively.

#### Algorithm 1: Depth Map Correspondence (DMCP)

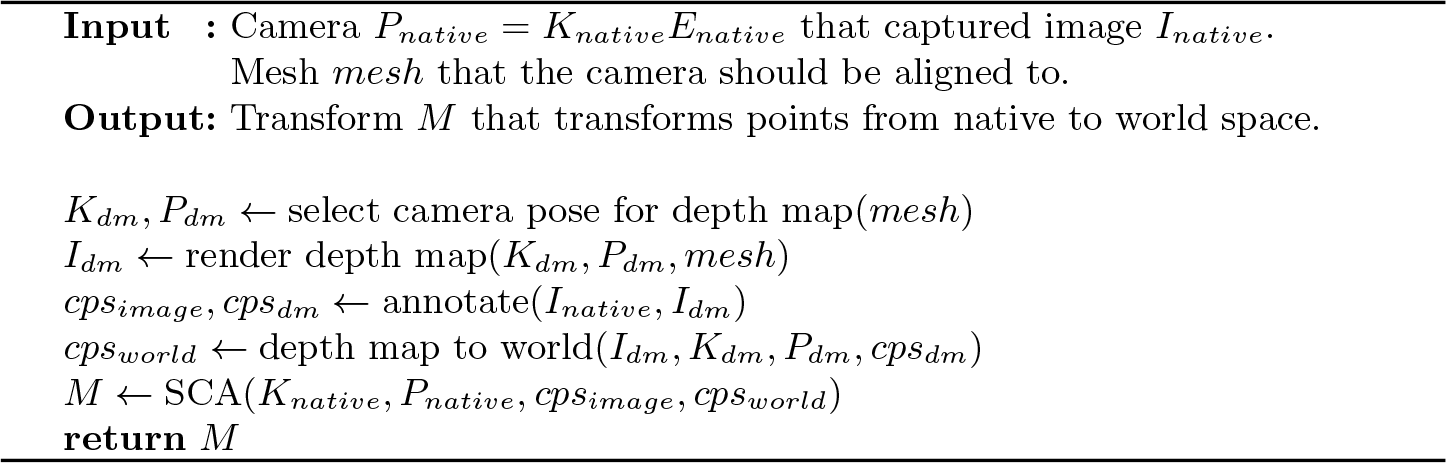

#### Algorithm 2: Sparse Camera Alignment (SCA)

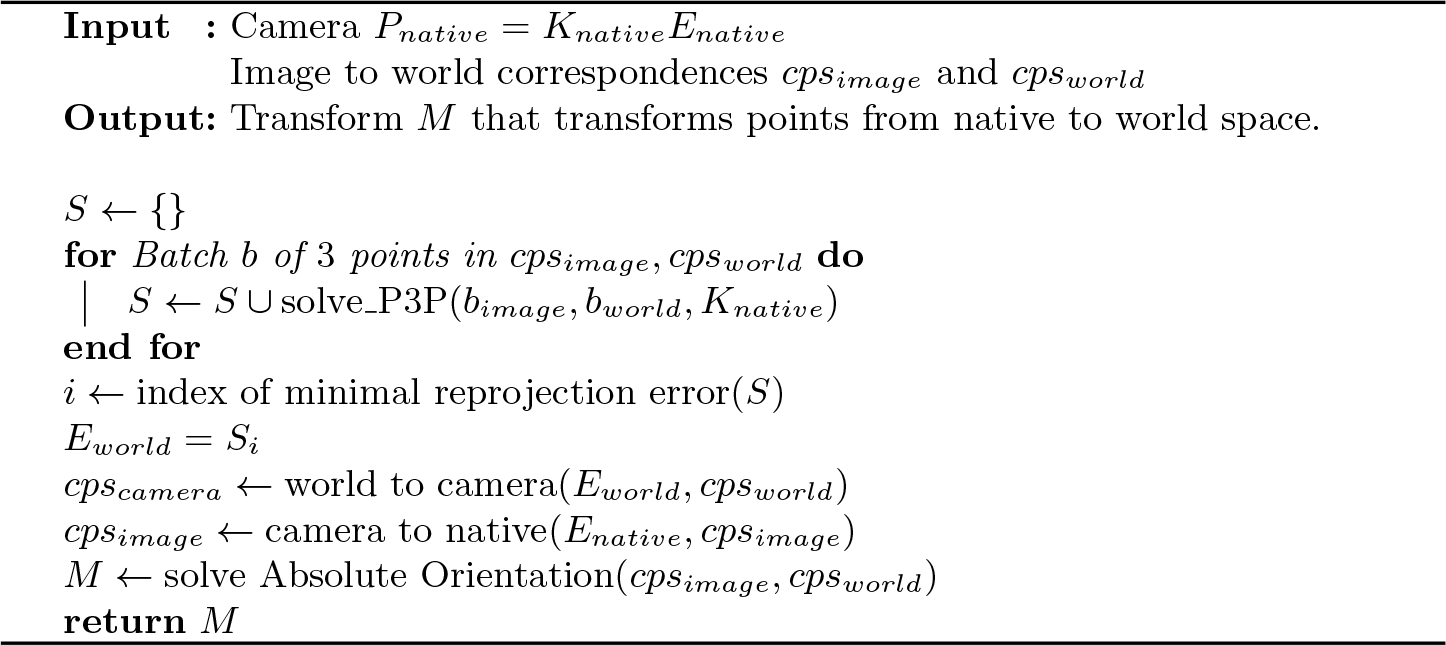

Let *a* ∈ *cps*_*dm*_ be the pixel coordinates of one of the corresponding points on *I*_*dm*_, such that *I*_*dm*_(*a*) is the depth. The respective point *A*_*camera*_ in the camera space of the depth camera can then be computed by

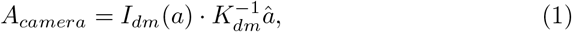

and further be transformed to world space with

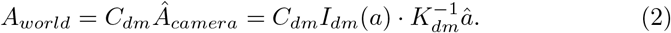

This is the computation which is performed in the *depth map to world* step in Alg. 1. The transformed set of correspondences in world space is denoted by *cps*_*world*_. Thus, *cps*_*image*_ and *cps*_*world*_ now represent correspondences from native pixel coordinates to world coordinates. Because correspondences are annotated and originate from a feature deficient scene, we assume their number to be close to the minimum number of 4 points necessary for the *Sparse Camera Alignment* step. Figure 2 shows example annotations.

### Sparse Camera Alignment

Suppose we have the native camera defined by its intrinsic camera matrix *K*_*native*_ and pinhole projection matrix *P*_*native*_ = *K*_*native*_*E*_*native*_. Additionally, we assume at least 4 corresponding points *cps*_*image*_ and *cps*_*world*_ generated in the annotation step. Sparse Camera Alignment is defined in two steps: 1) camera pose estimation and 2) transformation estimation.

#### Camera Pose Estimation

The position of the camera in the world is estimated using image to world correspondences. Since only the pose changes, the intrinsics *K*_*world*_ are the same as *K*_*native*_. This problem is known as the Perspective-*n*-Point problem. The lowest number of points with which camera pose can be estimated is 4 points. To reliably work on only 4 points, we build on an existing solution [10] of the Perspective-Three-Point Problem (P3P) implementated in [4]. P3P requires exactly three correspondences and gives up to 4 distinct valid solutions in the form of extrinsic matrices *E*_*i*_. The solutions are collected for all combinations of three points in the correspondence set. The fourth point (not input into P3P) is used to determine the correct solution among the *E*_*i*_. A point *A*_*world*_ ∈ *cps*_*world*_ can be reprojected into the image via

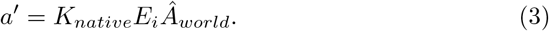

As every solution *E*_*i*_ is generated from 3 points, there is at least one point in *cps*_*world*_ that is not used to generate a solution. Thus, the solution that generalizes best can be selected as the solution where the total reprojection error

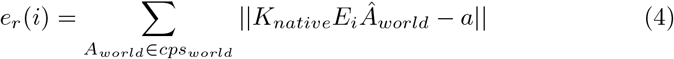

is minimized. Here, *a* ∈ *cps*_*image*_ is the point which corresponds to the projection *a*^*′*^ of the current point *A*_*world*_ for which the term is evaluated.

The final estimated extrinsic matrix and the solution of the camera pose estimation step is therefore

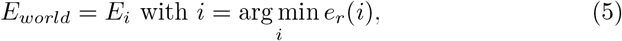

representing the estimated pose with respect to the world space.

#### Transformation Estimation

In this step, the transformation that transforms points in native space to world space is computed from the native camera *P*_*native*_ = *K*_*native*_*E*_*native*_, the estimated world camera *P*_*world*_ = *K*_*native*_*E*_*world*_ (note *K*_*native*_ = *K*_*world*_) and the annotated world points *cps*_*world*_.

First, each world point *A*_*world*_ ∈ *cps*_*world*_ is transformed moved to native space

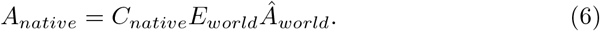

In this way, the coordinates of each world point in native space are known. The registering transformation can now be defined by solving the absolute orientation problem. Here, Umeyama’s method [21] as implemented in [4] is used. The estimated transformation *M* computes

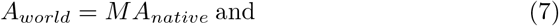

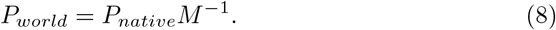

Thus, *M* solves the pose estimation problem for all cameras and points in native space. Alternatively, *M* could also be computed directly using the inversion *M* = *C*_*world*_*E*_*native*_. We use Umeyama’s method in our implementation since we found it leads to slightly more robust results.

### 2.2 Middlebury computational experiments

We demonstrate the generality of the *DMCP* algorithm to align any type of image and point-cloud/mesh using a pre-existing dataset with established groundtruth camera pose.

The *Art room 1* scene of the Middlebury dataset[19] is used to demonstrate *DMCP* ‘s generality. Each Middlebury scene consists of two rectified images and respective disparity maps, which are used to generate 3D point clouds (see Supplementary Information (SI)). The goal of this experiment is to find the original position of the camera. To simulate a native space that differs from world space, the native pinhole projection matrix is transformed with a random rotation and translation.

While *DMCP* requires a minimum of 4 points, we studied the effect of an increasing amount of annotated points. The accuracy of pose estimation with 4, 5, 6, 7, 8 and 16 correspondences is investigated with one trial per number of points.

### 2.3 The *Ushichka* dataset

The *Ushichka* dataset [2] consists of multiple nights of multi-sensor data capturing the flight behaviour of echolocating bats in the same recording volume. On each night, three uncooled thermal cameras (TeAx ThermalCapture 2.0, TeAx GmbH,Germany. Specifications: 640x512 pixels, 9mm focal length) were placed in similar locations in the cave across all nights. The DLT coefficients of each camera in the array was calibrated using [20] and 3D points calculated with [7]. The intrinsic and extrinsic camera parameters in native space are thus known. A high-resolution LiDAR scan of the cave was performed on 18th August 2018 (Stonex X300L scanner, ≤ 6 mm resolution). Details of the LiDAR scan and processing are described in [9]. The generated point cloud was downsampled to a centimetre level mesh for ease of handling. Only the relevant portion of the mesh corresponding to the recording volume is used in this work.

### 2.4. *Ushichka* computational experiments

#### Pose comparison and 2D reprojection errors

The nights between 21st July to 19th August 2018 of *Ushichka* are used for the experiment. The cameras in the dataset are placed at unique locations on each night, though their broad positions in the cave are similar across nights. Unlike the Middlebury dataset with one camera, here we have three cameras to calculate transformations from. Each thermal camera recorded a different view of the cave volume. There is a noticable variation in the thermal contrast based on camera location, recording time of night, and session date. The inter-camera variation thus affects the ability to find reliable correspondences between thermal image and depth map. The effect of camera view on pose estimation are studied. Corresponding points from each of the three cameras are used to generate transformations independently for each recording night. The transformations are then used to transform all cameras of the scene and compare their camera centers. For each camera and session, the variation in estimated position is quantified by calculating the Euclidean distances of pairs between the position estimates. The 2D reprojection error is compared between cameras on the same night, and across nights.

#### Microphone and cave point alignment

To verify the 3D alignment of the camera-LiDAR coordinate systems we compare the positions of microphones and other cave points after conversion to the LiDAR coordinate system. The multi-channel audio array consists of 4-18 small microphones that are placed on the cave wall. Microphone positions are ‘pointed’ using a hot incandescent bulb. The bulb’s 2D positions at each image are used to generate the 3D coordinates in native space. Additionally, up to 4 points on the cave (corners of a rectangular memorial plaque) are also pointed out using the bulb. Microphone and wall 3D points are then converted to LiDAR space using the session and camera specific transformations. A perfect alignment of microphone and LiDAR mesh means the microphone/cave points lie on the mesh surface. Alignment error is quantified by computing the distance to the nearest-mesh point for each transformed mic and wall point.

## 3 Results

### 3.1 Middlebury computational experiments

The Middlebury reprojection errors overall are low. Except for one outlier they all stay below 10*px* (see SI). From visual inspection, the reprojection error slightly increases with the amount of annotated correspondences (from 3 to 10 pixels across 4 to 16 annotated points). Also an outlier is seen at 7 annotations (50*px* error), indicating a false correspondence. Qualitatively, the camera pose shows an estimation error between 11-27 cm across 4-16 annotations (Figure 3 and see SI for details). While it may seem that more annotations could provide a better pose estimate, more annotations also increase the chances of mis-annotation. In general the Middlebury experiment validates the algorithm and demonstrates that a reasonable pose is found. Further it shows that the quality of the correspondences is more important than its quantity.

**Fig. 3:**
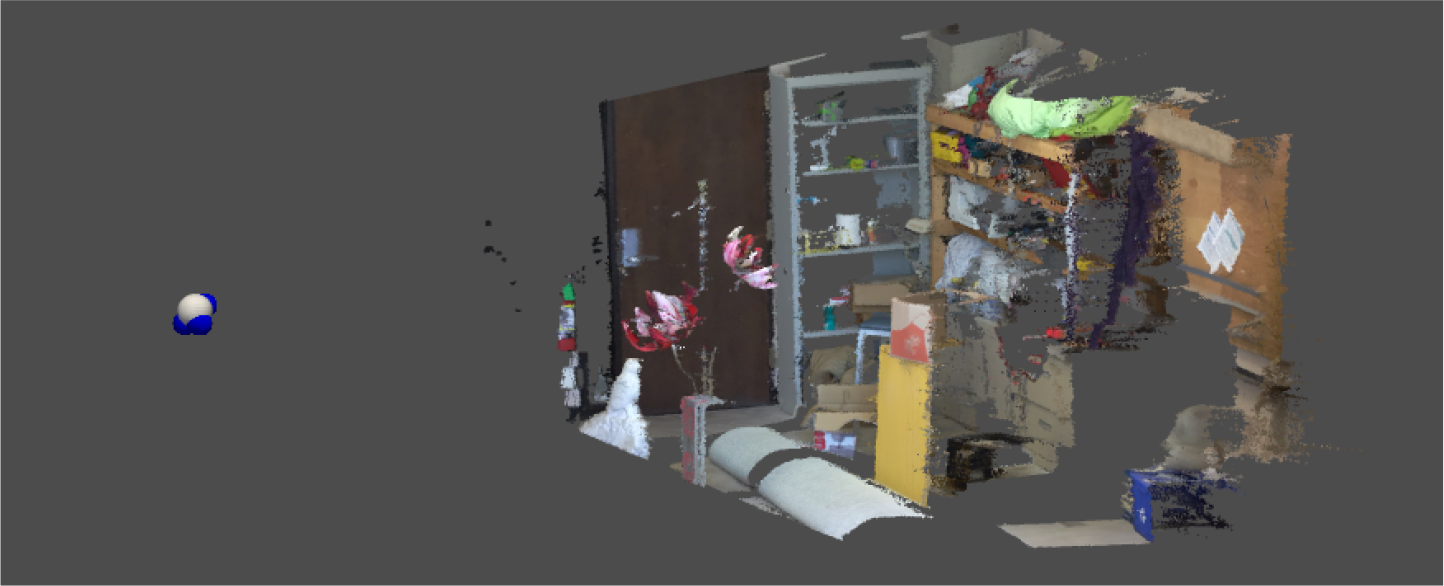
Estimated camera positions relative to the scene. *6* positions are shown as blue spheres (not all visible), the original position is a white sphere.

### 3.2 Ushichka Computational Experiments

#### 2D reprojection errors and camera pose variation

We calculate the mean reprojection error across cameras for the 4th point on each recording night. The errors in pixels are (from 2018-07-21 to 2018-08-19): 3.97, 8.30, 23.74, 6.71, 6.61, 3.81 and 21.81. While there are two nights with high errors (2018-07-28 and 2018-08-19), 5 of 7 nights show ≤ 10 pixels and 2 nights even show ≤ 5 pixels error. All transformations are estimated using 4 correspondences. Figure 5 shows the distances between the estimated poses for each camera. The interpretation is that if two transformations agree on a similar position for a camera, that position likely is correct. The minimum distance captures this concept. The maximum distance captures the concept that if the transformations are wrong, they move cameras to an arbitrary position which increases the maximum distance between estimated camera positions.

Interestingly, recording *08-14* has a low mean reprojection error, while large minimum camera distance and even larger maximum distance. The explanation is that one transformation was almost correct while the other two transformations are wrong. We know this as the last transformation produces camera positions in an area where we know them to be placed in reality. When comparing an image from *2018-08-14* (see SI) with an image from *2018-08-17* (Figure 2a) where all transformations performed reasonably, it becomes clear that the image quality of recording *2018-08-14* makes it much harder to annotate correctly. The camera positions of *2018-08-17* are visualized in Figure 4.

**Fig. 4:**
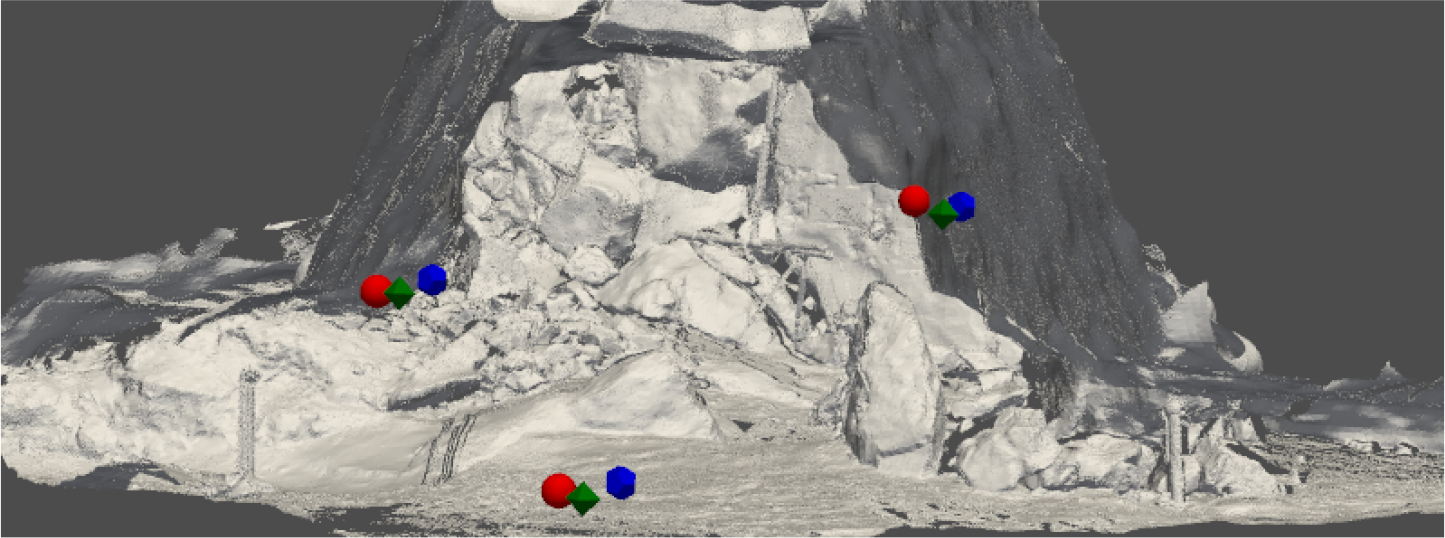
Estimated camera positions using transform estimated from camera 1 (red sphere), 2 (green octahedron) and 3 (blue dodecahedron) for 2018-08-17.

#### Microphone and cave point alignment

Ideal camera-LiDAR alignment should result in 0 distance between camera-triangulated points and nearest mesh points. We observe a range of alignment distances (Figure 6) with a median of 1 m (95%ile: 0.008-5.61 m, *N* =261 points). Some nights showed good alignment. For 3 sessions, the maximum nearest-point distance is ≤ 0.5 m (2018-07-21, 2018-07-25, 2018-08-17: 0.38, 0.32, 0.45 m). It is important to caution that a lower nearest-point distance implies good fit of camera triangulated points with the mesh, but does not automatically imply a good alignment. For instance, in some cases the transformation leads to the points being close to the mesh but not in the expected location (e.g. higher or lower than expected in the cave).

In general, the experiment shows that the *DMCP* framework empowers a user to register calibrated cameras to a mesh, even in difficult scenes. Especially the difficult images in the *Ushichka* dataset that did not result in a correct transformation can be estimated simply by trying different correspondences and taking more time to find correct ones.

**Fig. 5:**
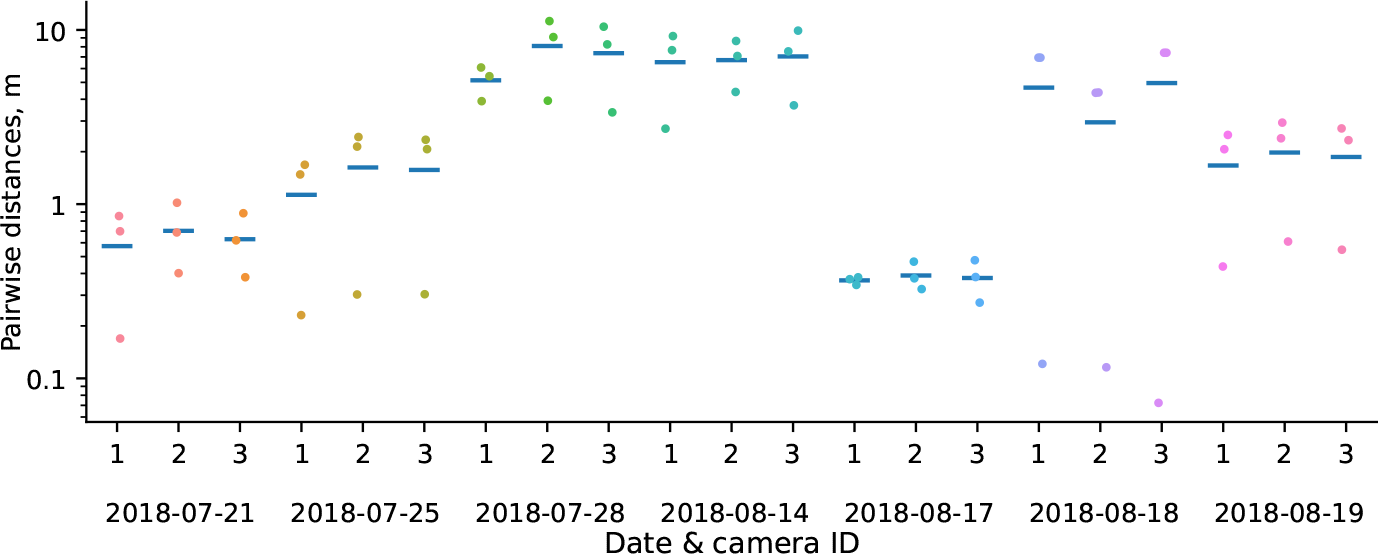
Quantifying pose estimate variation for each camera based on the source transformation. Each camera has three pose estimates produced by the transform generated from each camera image. The pairwise distances for the three possible pairs are plotted on a *log*_10_ scale. A consistent pose estimate will show pairwise distances close to zero. Camera pose variation within 1 metre distance is seen when reliable correspondence annotations are possible (e.g. 2018-08-17 and 2018-07-21). High variation in pose estimate doe not necessarily imply that all of the estimated poses are wrong. The mean for each camera is indicated with a horizontal line.

**Fig. 6:**
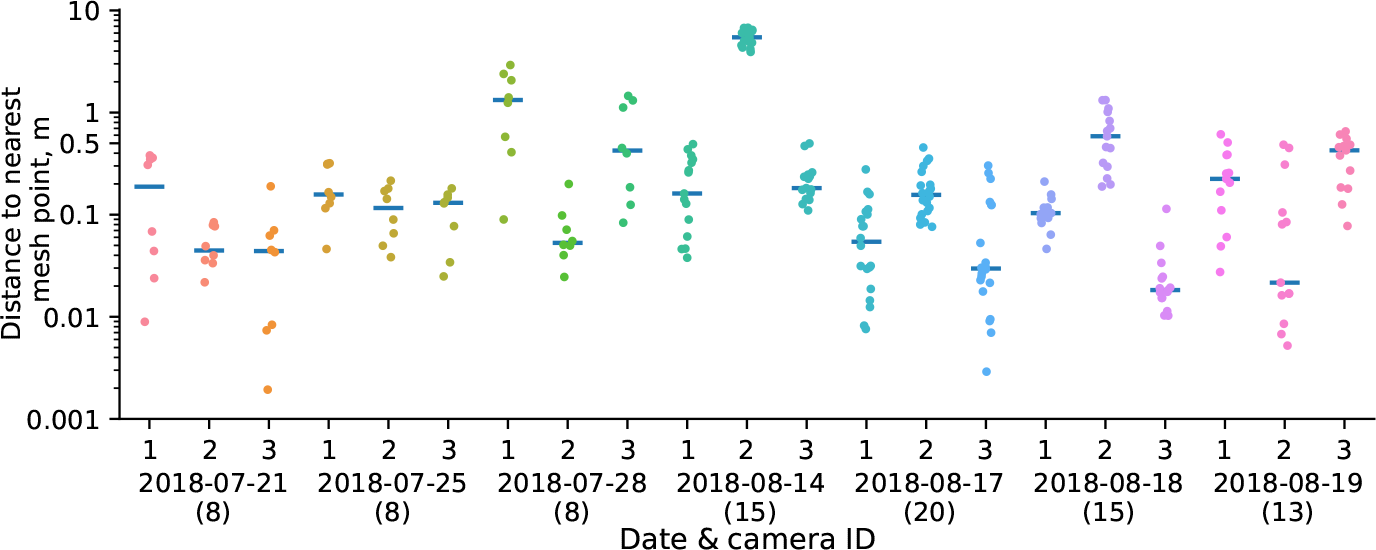
Quantifying alignment error: distance between transformed microphone and cave points and the nearest LiDAR mesh point. Y-axis in *log*_10_ scale. X-axis shows camera ID used to generate correspondences, session date, and number of microphone and wall points used to quantify alignment error. Horizontal lines are medians. Most of the points are within 1 m (90th percentile) of a mesh point. A clear effect of camera placement and annotation ease is seen, for example Camera 2 on 14th August with transformed points that do not lie in the mesh itself.

## 4 Discussion

Sensor fusion is of particular importance given the variety of commercially available sensors, and physical conditions under which animal behaviourists deploy these sensors. To aid in thermal camera-LiDAR alignment we developed the DCMP algorithm. DMCP estimates camera pose using user-assisted correspondences between thermal images and LiDAR depth maps. We highlight the success of the method despite the limited thermal camera resolution (640x512 pixels), challenging image conditions (2a, also see SI), along with the low number (4 points) of correspondences required from the user’s side. Finding reliable correspondences between images is challenging even during manual annotation because of the low thermal contrast across the cave, varying camera sensitivities, and changing thermal conditions across each night. To improve image quality, we found it sometimes useful to generate median filtered images derived from multiple frames across the night. Given the abundance of interactive 3D visualisation tools, one may question our choice of using a depth map for correspondences between the LiDAR and thermal image. Our experience shows that clicking on a depth map which resembles the thermal image is far more intuitive, and resultes in more reliable correspondences. Clicking on corresponding points in a 3D interactive session of the scene is much more challenging and time consuming than expected.

DMCP is a user-assisted algorithm, meaning that poor input correspondences will result in a poor pose estimation. In particular, point annotations made close to edges in the depth map can tip the balance between a good and bad pose estimation. As observed in the Middlebury experiments (section 3.1), a few reliable correspondences are better than many poor ones. The intrinsic and extrinsic parameters of the experimental cameras are assumed to be known. Errors in camera parameter estimation (the effect of using various calibration workflows) will also influence DMCP’s accuracy. A last but important point to note is to ensure the consistency of camera and LiDAR coordinate systems (left/right hand coordinate systems, see SI). The various sources of error in *DMCP* may mean that resulting transformation matrices are sensible, but may not fit as closely as required. Here, we found the iterative-closest-point (ICP) algorithm [3] of great use (data not shown). When the user has camera 3D coordinates of points on the LiDAR scene (as in 3.2), ICP can help improve the overall fit of thermal camera and LiDAR data.

Even though DMCP is developed to solve thermal camera-LiDAR alignment, we stress that it is generalisable to any kind of camera-pointcloud or camera-mesh data. We, in fact, tested *DMCP* using the camera-point cloud Middlebury dataset, and established the correctness of the algorithm. Other potential uses of DMCP are in generating pose estimation for aligning meshes generated from any 3D scanning device or point-clouds generated from structure-from-motion type workflows.

## 5 Conclusions

We found it surprising that there currently does not seem to be an automatic solution which works to align multiple thermal cameras calibrated relative to each other to a mesh - finding correspondences between the thermal images to reliably obtain 3D points on the mesh seems to be extremely challenging in scenes with low temperature variation . We believe this would be an interesting avenue for future work.

The current formulation of DMCP estimates transforms from single cameras independently. Future work could generalize the algorithm to multiple cameras. This could be achieved by estimating a consensus transform using point correspondences from all cameras together. The combined approach transforms may result in a more robust transform estimate by removing outliers that do not satisfy the relative camera pose constraint. *DMCP* is a good building block for a completely automated method, which is likely to scale well as the number of cameras and recording sessions increase. We share the code and relevant raw data to encourage further research in this direction (see SI). Automation of two portions of the workflow require further development: 1) generation of corresponding points, and 2) generation of depth maps. To automatically find corresponding points, future work could look into feature detectors optimised from thermal images, such as phase-congruency [12,5,8], or machine learning approaches using larger training datasets. Instead of a single depth map used to obtain corresponding points, an automated method could generate multiple views of the mesh with varying numbers of corresponding points from each depth map. The corresponding points across depth maps could then be pooled to obtain a robust pose estimate.

## Supporting information

Supplementation Information to main MS

