## Supplementation Information to main MS for "A target-free approach to estimate thermal camera pose in LiDAR scenes of feature-deficient environments"

### 1 Testing DMCP with Middlebury dataset

#### 1.1 Generating point clouds from disparity maps

Also, the camera's intrinsics $K_{native}$, focal length $f$, disparity offset $o$ and baseline $b$ are known. For this experiment, the first rectified image and disparity map of the scene *artroom1* was taken. The disparity map $D'$ is converted into a depth map $D$. For each non-zero disparity at pixel $a$, the associated depth

$$D(a) = f \cdot \frac{b}{D'(a) + o} \tag{1}$$

is computed. Every $D(a)$ is converted into a point cloud by computing the world point

$$A_{world} = D(a) \cdot K_{native}^{-1} \hat{a}. \tag{2}$$

This point cloud

$$PC = \{A_{world}(a) \mid a \text{ is pixel with non-zero disparity in } D'\} \tag{3}$$

is now considered the world space.

#### 1.2 Results

| Number of annotated points | 4 | 5 | 6 | 7 | 8 | 16 |
|---|---|---|---|---|---|---|
| Distance from groundtruth position, m | 0.27 | 0.17 | 0.11 | 0.25 | 0.13 | 0.20 |

Table 1: Error in camera pose estimation for the Middlebury dataset

We estimate camera pose error by calculating the euclidean distance between estimated and ground truth position. As we see in Table 1, the positions were estimated fairly close to ground truth with distances between $11 - 27cm$. Considering the distance of the Middlebury camera from the scene we believe the pose estimation error observed is relatively low. From visual inspection (and given the limited sample size), it is difficult to say if there is a correlation between number

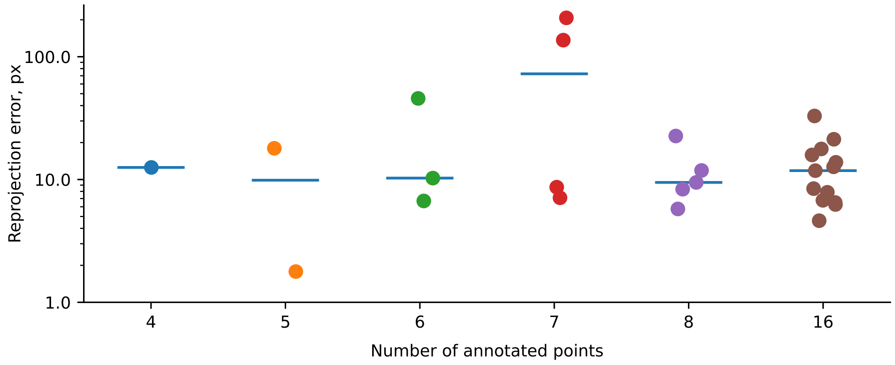


Fig. 1: Reprojection error of annotated Middlebury points. Horizontal line shows median. The non-zero $N_{annotated} - 3$ reprojection errors are plotted as points. Y-axis is in $\log_{10}$ scale. The results qualitatively illustrate the importance of good quality annotations, rather than their number.

of annotations and pose estimation error. Our general observations hint, that the *quality* of annotations is more important than the quantity.

For each run of DMCP we obtain one estimated pose, and a number of non-zero 2D reprojection errors. The reprojection errors of three out of $N$ annotated points is typically very close to zero. These are the three points actually used to solve for the camera position with P3P. The reprojection error of the other $N-3$ points is however more informative of estimation error. A low reprojection error of these $N-3$ points implies a good camera pose estimate and consistency between all thermal-LiDAR annotations. We see from Figure 1 that the reprojection error is generally low with a median of 11 pixels (all non-zero error points pooled). We see a few outliers (e.g. at 7 annotated points) with very high reprojection errors of 136 and 207 pixels. These high reprojection errors are likely due to bad annotations.

### 2 DMCP on Ushichka: challenges and details

#### 2.1 Difficult thermal images: example

The variation in camera sensitivity, weather and daily thermal conditions affect the clarity of the thermal image that needs to be annotated. Some nights provide clear images where cave features can be comparatively easily distinguished and annotated, (e.g. see Fig. 2a Main Text). Other nights however are much more difficult to annotate. *2018-08-14*, shown in Figure 2 for instance is particularly challenging to annotate.

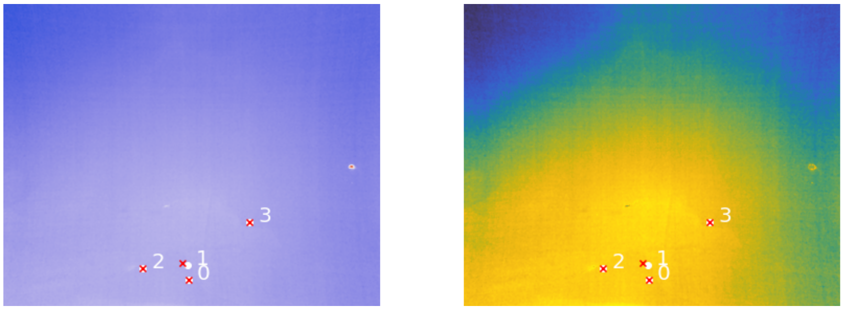


(a) coolwarm colormap

(b) gouldian colormap, high colors (colors $>$ mean $+ 2 \cdot$ std) are remapped to mean

Fig. 2: Thermal image of night 2018-08-14. Annotated thermal points are shown in white, the depth map reprojections are indicated as a red x. Figure 2a shows the same color palette as 2018-08-17 for consistency (see Main Text Figure 2a). It is possible to improve the visualisation a little bit by optimizing it for the image specifically (Figure 2b)

#### 2.2 Consistent camera coordinate systems

This is a detail specific to the Ushichka dataset, but perhaps likely to be encountered by other experimenters too. We initially faced problems aligning the thermal camera and LiDAR coordinate systems despite having good quality user-annotations. On further inspection we discovered, the problem with alignment arose from the fact that the LiDAR coordinate system was right-handed, while the DLT calibration and camera triangulation implemented in [1,2] used a left-handed coordinate system.

The thermal camera and LiDAR could be aligned only after explicitly accounting for this discrepancy in handedness. Specifically, every point $A_{native}$ in native space that corresponds to a point $A_{world}$ in world space needs to be transformed to the same point $A_{camera}$ in camera space by the extrinsic matrices $E_{native}$ and $E_{world}$ respectively.

### 3 Code and raw data for replication and further development

*Thermal camera and LiDAR data* To aid in the further development of thermal camera-LiDAR pose estimation 'in-the-wild', we share the LiDAR-camera data on each of the 7 recording nights in the *Ushichka* dataset. The dataset is uploaded with the DOI: 10.5281/zenodo.6620671

*DMCP code* The version of the DMCP code used to generate the results of this paper have the DOI: 10.5281/zenodo.6621388. The codebase is released under an MIT software license, enabling further development by the community.
